# Mobile elements habouring heavy metal and bacitracin resistance cassettes are common among *Listeria monocytogenes* persisting on dairy farms

**DOI:** 10.1101/2021.03.15.435412

**Authors:** Hanna Castro, Francois Douillard, Hannu Korkeala, Miia Lindström

**Author notes:** **Corresponding author:** Hanna K. Castro.

## Abstract

*Listeria monocytogenes* is a food-borne pathogen and a resilient environmental saprophyte. Dairy farms are a reservoir of *L. monocytogenes* and strains can persist on farms for years. Here, we sequenced the genomes of 250 *L. monocytogenes* isolates to investigate the persistence and mobile genetic elements of *Listeria* inhabiting dairy farms. We performed a SNP-based phylogenomic analysis to identify 14 monophyletic clades of *L. monocytogenes* that persistent on the farms for ≥ 6 months. We found that prophages and other mobile genetic elements were on average more numerous among isolates in persistent than nonpersistent clades, and demonstrated that resistance genes against bacitracin, arsenic and cadmium were significantly more prevalent among isolates in persistent than nonpersistent clades. We identified a diversity of mobile elements among the 250 farm isolates, including three novel plasmids, three novel transposons and a novel prophage harbouring cadmium resistance genes. Several of the mobile elements we identified in *Listeria* were identical to the mobile elements of *Enterococci*, indicative of recent transfer between these genera. Finally, we demonstrated that the CRISPR-*cas* IIa system and a type II restriction-modification system were negatively associated with persistence on farms. Our findings suggest that mobile elements support the persistence of *L. monocytogenes* on dairy farms and that *L. monocytogenes* inhabiting the agroecosystem is a potential reservoir of mobile elements that may spread to the food industry.

**Importance:** Animal derived raw materials are an important source of *L. monocytogenes* for the food industry. Knowledge of the factors contributing to the pathogen’s transmission and persistence on farms are essential for designing effective strategies against the spread of the pathogen from farm to fork. An increasing body of evidence suggests that mobile genetic elements support the adaptation and persistence of *L. monocytogenes* in the food industry, as these elements contribute to the dissemination of genes encoding favourable phenotypes, such as resilience against biocides and thermal stress. Understanding the role of farms as a potential reservoir of these elements is needed for managing the transmission of mobile elements across the food chain. Because *L. monocytogenes* coinhabits the farm ecosystem with a diversity of other bacterial species, it is important to assess the degree to which genetic elements are exchanged between *Listeria* and other species, as such exchanges may contribute to rise the novel resistance phenotypes.

## Introduction

*Listeria monocytogenes* leads a double life: in one it is a potentially lethal, zoonotic foodborne pathogen and in the other, a ubiquitous environmental saprophyte (Gray et al., 2006). Agroecosystems provide a favourable habitat for *L. monocytogenes* and the pathogen is especially prevalent on dairy farms (Nightingale et al., 2004; Esteban et al., 2009). *L. monocytogenes* strains can inhabit dairy farms for years and be widely distributed in the farm environment, leading to the frequent contamination of milk (Ho et al., 2007; Castro et al., 2018). Raw milk and animals destined for slaughter are a major contamination source for the food industry (Samelis & Metaxopoulos 1999; Fox et al., 2009; Hellström et al., 2010). Knowledge of the pathogens ecology on farms is essential for controlling the spread of *L. monocytogenes* from farms to the food industry.

*L. monocytogenes* is extremely resilient and can tolerate various stresses used to control the pathogen in the food industry (Lundén et al., 2003; Aarnisalo et al., 2007). These phenotypic traits enable *L. monocytogenes* to survive in food processing environments for years, a phenomenon known as persistence (Lundén et al., 2000; Keto-Timonen et al., 2007; Stasiewicz et al., 2015; Pasquali et al., 2018; Hurley et al., 2019). Mobile genetic elements are common among *L. monocytogenes* isolates from food processing environments (Harvey & Gilmour, 2001; Pasquali et al., 2018; Hurley et al., 2019) and may harbour genes mediating tolerance to heat shock (Pöntinen et al., 2017), salt and acid stress (Naditz et al., 2019; Hingston et al., 2019) and biocides (Müller et al., 2013; Meier et al., 2017). These findings led us to the hypothesis that mobile genetic elements play a key role in the environmental adaptation and persistence of *L. monocytogenes*.

Although dairy farms are considered a reservoir of *L. monocytogenes* (Nightingale et al., 2004), and are known to harbour hypervirulent strains (Maury et al., 2019), the era of next generation sequencing has witnessed very few efforts to illuminate the pathogen’s ecology in the farm environment. How *L. monocytogenes* adapts to life in the farm ecosystem, and to what extent the farm environment acts as source of mobile genetic elements for *L. monocytogenes* persisting in food processing environments, are key issues to explore. Such insights would be instrumental in developing novel strategies to reduce contamination on farms and in the raw materials delivered to the food industry.

Here, we sequenced the genomes of 250 *L. monocytogenes* isolates obtained from three Finnish dairy farms during 2013–2016 (Castro et al., 2018) to investigate the persistence and mobile genomic elements of *L. monocytogenes* in the farm environment. We performed a SNP-based phylogenomic analysis to group the isolates into persistent and nonpersistent clades and identified plasmids and chromosomal mobile elements among the 250 genomes. We found that prophages and other mobile genetic elements were on average more abundant among isolates in persistent than nonpersistent clades, and that a significantly higher portion of isolates in persistent than nonpersistent clades harboured genes against bacitracin, arsenic and cadmium. Finally, we explored genome wide association between gene content and persistence. We found a negative association between persistence and three putative defence systems against invading prophages and plasmids (Garneau et al, 2010; Lee et al., 2012). Taken together, our findings suggest that prophages and mobile genetic elements confer an ecological advantage for persistence on farms, and that *L. monocytogenes* inhabiting the farm environment constitutes a reservoir of diverse mobile genetic elements that may spread upstream the food chain.

## Results

### Persisting clades of *L. monocytogenes* were detected on all three farms

Whole genome sequencing and subsequent *in silico* subtyping of 250 *Listeria monocytogenes* isolates, collected from three Finnish dairy farms during 2013–2016 (Castro et al., 2018), yielded 25 unique multi-locus sequence types (STs) (Fig. 1a, see Supplementary Data S1). The most frequently detected subtype was ST20, which represented 28% of all sequenced isolates. In this study, persistent clades of *L. monocytogenes* were defined as monophyletic clades of isolates with pair-wise distances (PWDs) fewer than 20 SNPs (Pightling et al., 2018) that were isolated from the same farm from ≥3 samples during ≥6 months. Clades that did not meet these criteria were classified as nonpersistent. We identified in total 14 persistent clades (Fig. 2, Table 1). Persistent clades represented 71% of all sequenced isolates and all persistent clades belonged to serogroup 1/2a. Clade C4 contained isolates from two different farms, suggesting that strains of *L. monocytogenes* can spread between farms faster than the rate of genomic diversification.

**Fig. 1.**
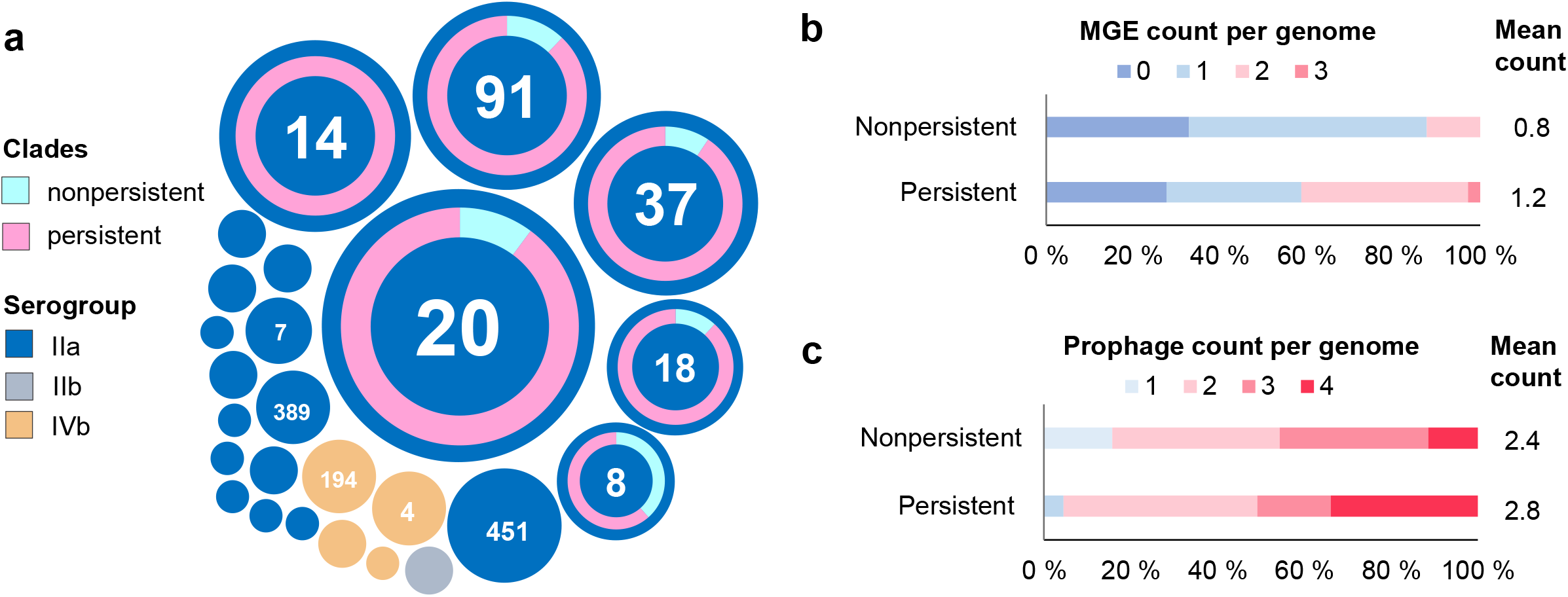
*L. monocytogenes* isolates in persistent clades contained on average more prophages and other mobile genetic elements (MGE) than isolates in nonpersistent clades. **a** *L. monocytogenes* isolates of this study represented 25 unique STs and persistent clades were detected among the six most prevalent STs. Each circle represents a unique ST and the area of the circle corresponds to the number of isolates. Doughnut charts illustrate the proportion of persistent clades (pink) and singleton isolates (aquamarine) for each STs in which persistent clades were detected. **b**, **c** Distribution of isolates by number of non-phage MGE (**b**) and prophages (**c**) per genome among isolates in persistent and nonpersistent clades. The average number of the elements per genome is also given.

**Fig. 2.**
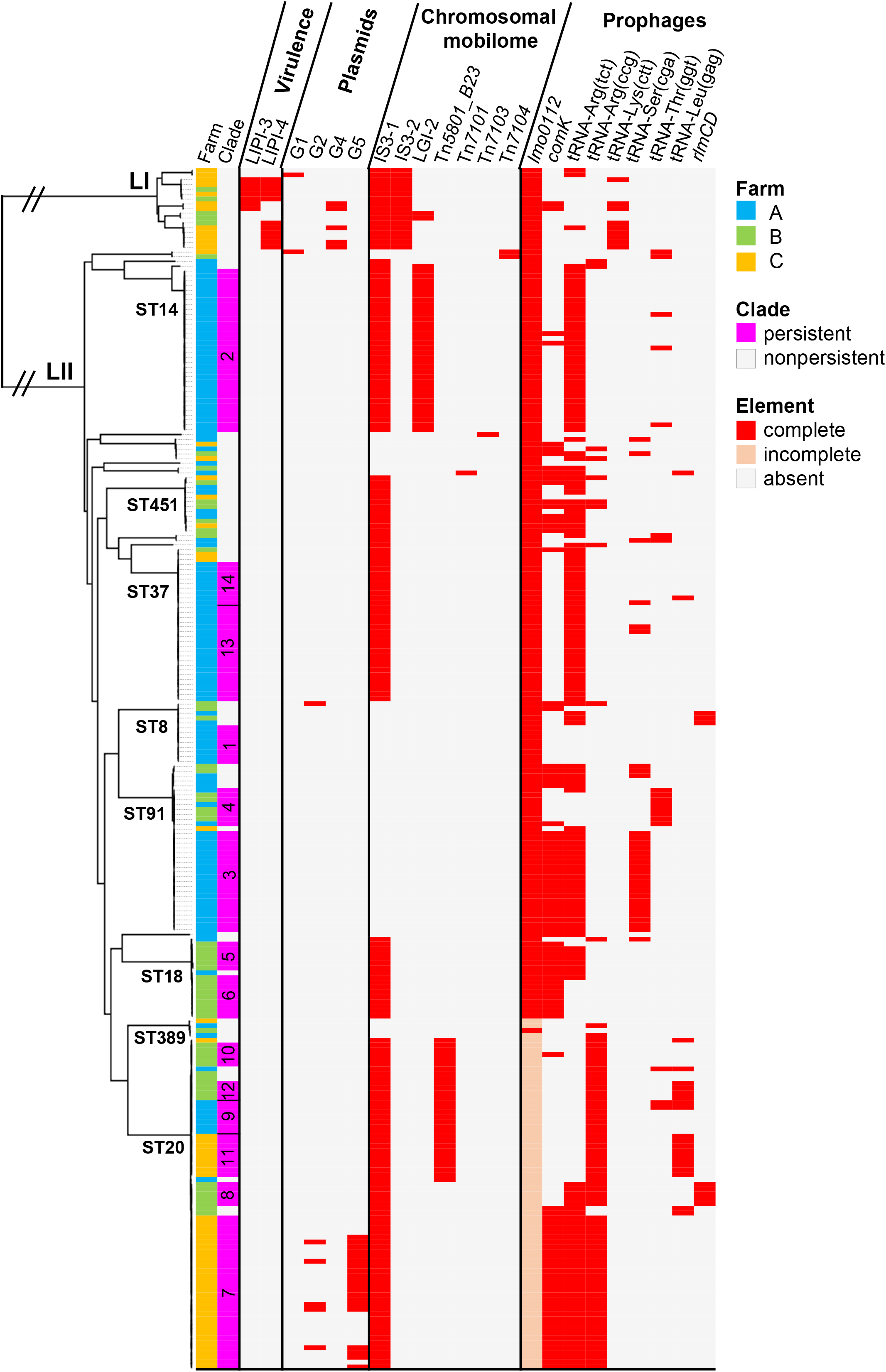
Phylogeny and genomic elements of 250 *L. monocytogenes* dairy farm isolates. The Lyve-SET 1.1.4f SNP-calling pipeline was used to generate an alignment file of the 250 genomes using *L. monocytogenes* EGD-e (NC_003210.1) as reference. Recombinant sites were removed from the alignment using Gubbins 3.0. Maximum likelihood phylogeny was inferred from concatenated SNP alignment files using PhyML 3.3. The tree was visualized using FigTree 1.4.4. Pathogenicity islands, plasmids, chromosomally located mobile elements and prophages were identified from the assemble and annotated draft genomes. The heatmap is restricted to genomic elements that were detected in this study. Persistent clade numbers corresponding to Table 1 are shown. Plasmids are categorized by phylogenetic group and prophages by insertion site. L: lineage; ST: multi-locus sequence type; LIPI: *Listeria* Pathogenicity Island; IS3: *Listeria* IS3-like element; LGI-2: *Listeria* Genomic Island 2.

**TABLE 1.**
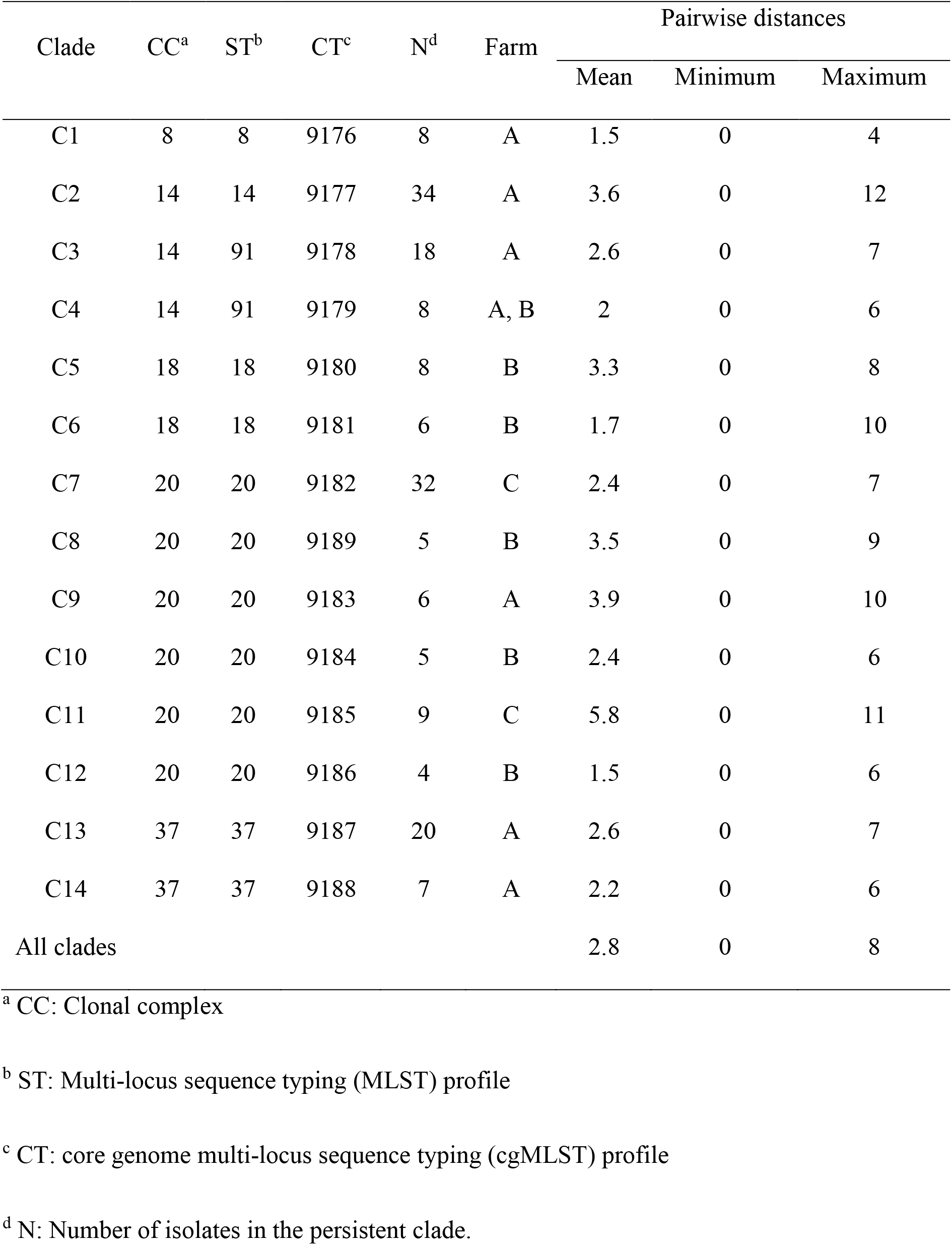
Pairwise distances within persistent clades of *L. monocytogenes* from dairy farms A – C.

| Clade | CC<sup>a</sup> | ST<sup>b</sup> | CT<sup>c</sup> | N<sup>d</sup> | Farm | Pairwise distances |  |  |
| --- | --- | --- | --- | --- | --- | --- | --- | --- |
|  |  |  |  |  |  | Mean | Minimum | Maximum |
| C1 | 8 | 8 | 9176 | 8 | A | 1.5 | 0 | 4 |
| C2 | 14 | 14 | 9177 | 34 | A | 3.6 | 0 | 12 |
| C3 | 14 | 91 | 9178 | 18 | A | 2.6 | 0 | 7 |
| C4 | 14 | 91 | 9179 | 8 | A, B | 2 | 0 | 6 |
| C5 | 18 | 18 | 9180 | 8 | B | 3.3 | 0 | 8 |
| C6 | 18 | 18 | 9181 | 6 | B | 1.7 | 0 | 10 |
| C7 | 20 | 20 | 9182 | 32 | C | 2.4 | 0 | 7 |
| C8 | 20 | 20 | 9189 | 5 | B | 3.5 | 0 | 9 |
| C9 | 20 | 20 | 9183 | 6 | A | 3.9 | 0 | 10 |
| C10 | 20 | 20 | 9184 | 5 | B | 2.4 | 0 | 6 |
| C11 | 20 | 20 | 9185 | 9 | C | 5.8 | 0 | 11 |
| C12 | 20 | 20 | 9186 | 4 | B | 1.5 | 0 | 6 |
| C13 | 37 | 37 | 9187 | 20 | A | 2.6 | 0 | 7 |
| C14 | 37 | 37 | 9188 | 7 | A | 2.2 | 0 | 6 |
| All clades |  |  |  |  |  | 2.8 | 0 | 8 |

<sup>a</sup> CC: Clonal complex
<sup>b</sup> ST: Multi-locus sequence typing (MLST) profile
<sup>c</sup> CT: core genome multi-locus sequence typing (cgMLST) profile
<sup>d</sup> N: Number of isolates in the persistent clade.

Pathogenicity islands associated with hypervirulence (LIPI-3, LIPI-4) were detected in 5% of the 250 isolates, none of which belonged to persistent clades. None of the 250 isolates harboured a premature stop codon within the *inlA* gene, which is associated with hypovirulence and is a common finding in *L. monocytogenes* form food processing environments (Maury et al., 2016). Indeed, the two STs most strictly associated with the food processing environments, namely 9 and 121 (Maury et al., 2016), were not detected in this study.

### Mobile genetic elements were on average more numerous among isolates in persistent than nonpersistent clades of *L. monocytogenes*

Overall, prophages and other mobile genetic elements were significantly more numerous (Independent Samples Median Test; *p*<.01) among isolates in persistent than nonpersistent clades (Fig. 1b-c). Resistance cassettes against cadmium and arsenic were detected in 20 and 15 % of isolates, respectively. Mobile elements harbouring resistance genes against arsenic and cadmium were significantly more prevalent among persistent clade isolates than singleton isolates (Fig. 3d). Surprisingly, 12% of all *L. monocytogenes* isolates, harboured putative bacitracin resistance cassette (Manson et al., 2004), located on the transposon Tn*5801*_B23. Other antimicrobial or biocide resistance genes were not detected in this study.

**Fig. 3.**
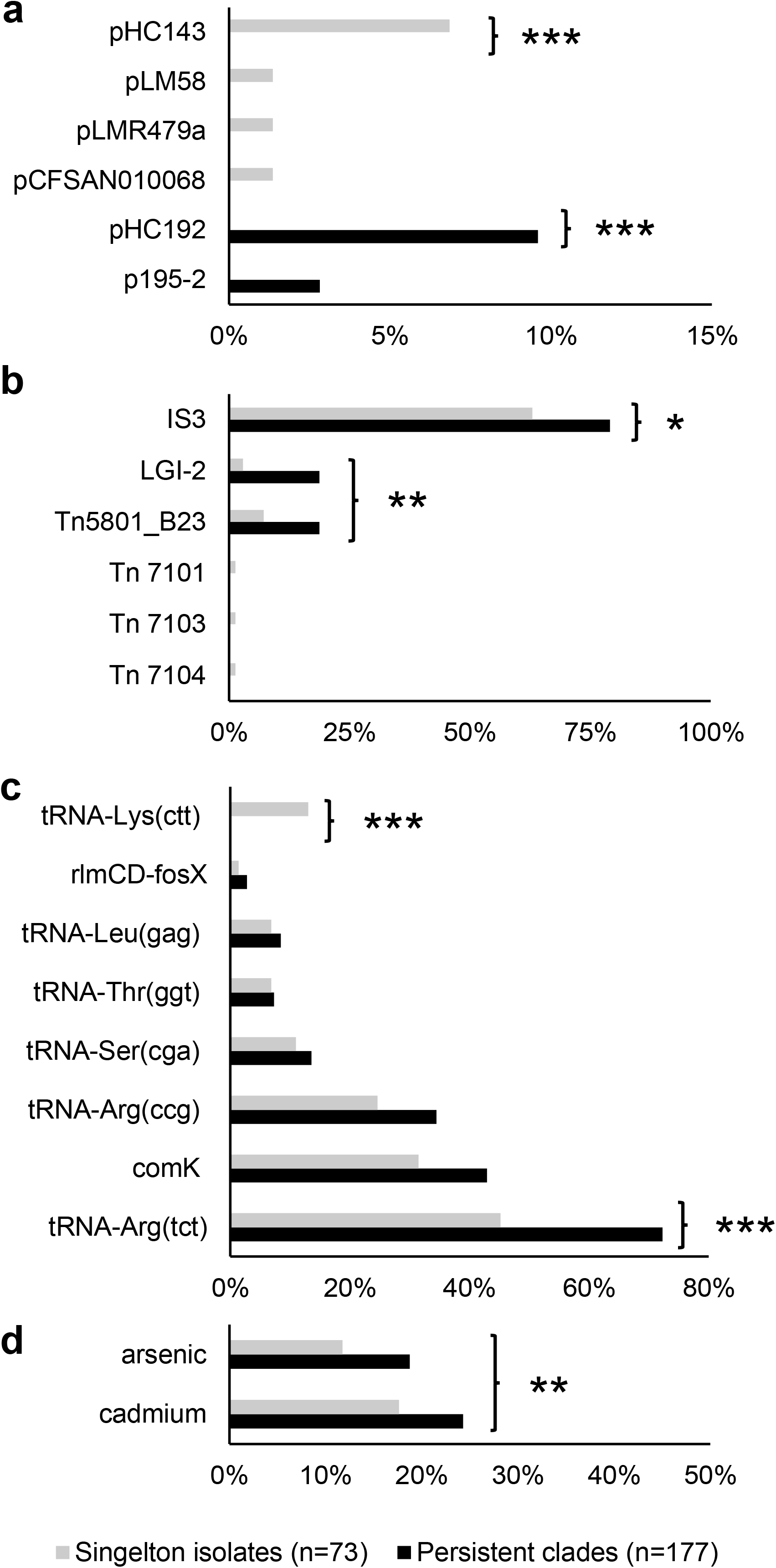
Occurrence of mobile genetic elements and heavy metal resistance genes among persistent clades and nonpersistent. Occurrence of plasmids (**a**), chromosomally located mobile elements (**b**), prophages (**C**) and cadmium and arsenic resistance genes (**d**) among isolates in persistent and nonpersistent clades. Significant differences between persistent clade isolates and singleton isolates are denoted by asterisks; *: *p*<.05; *: *p*<.05; ***: *p*<.001. IS3: *Listeria* IS3-like transposon; LGI-2: *Listeria* Genomic Island 2. Prophages are categorized by insertion site.

### Dairy farm isolates of *L. monocytogenes* harboured plasmids that are common in the food industry and three novel plasmids

Plasmids were detected among 10% of *L. monocytogenes* isolates in persistent clades and 11% singleton isolates. We detected three previously identified plasmids (pCFSAN010068, pLM58, pLMR479a), and three novel plasmids, labelled pHC143, pHC192 and pHC195-2 (Fig. 3a, see Supplementary Data S1). These plasmids were 55.5 – 86.7 Kb in size, except for pHC192, which was only 4.6 Kb. A Maximum-Likelihood phylogenetic analysis based on RepA grouped the five large plasmids into the plasmid groups G1, G2 and G4 (Kuenne et al., 2010; Schmitz-Esser et al., 2021), which appear to be specific to the genus *Listeria* (Fig. 4a).

**Fig. 4.**
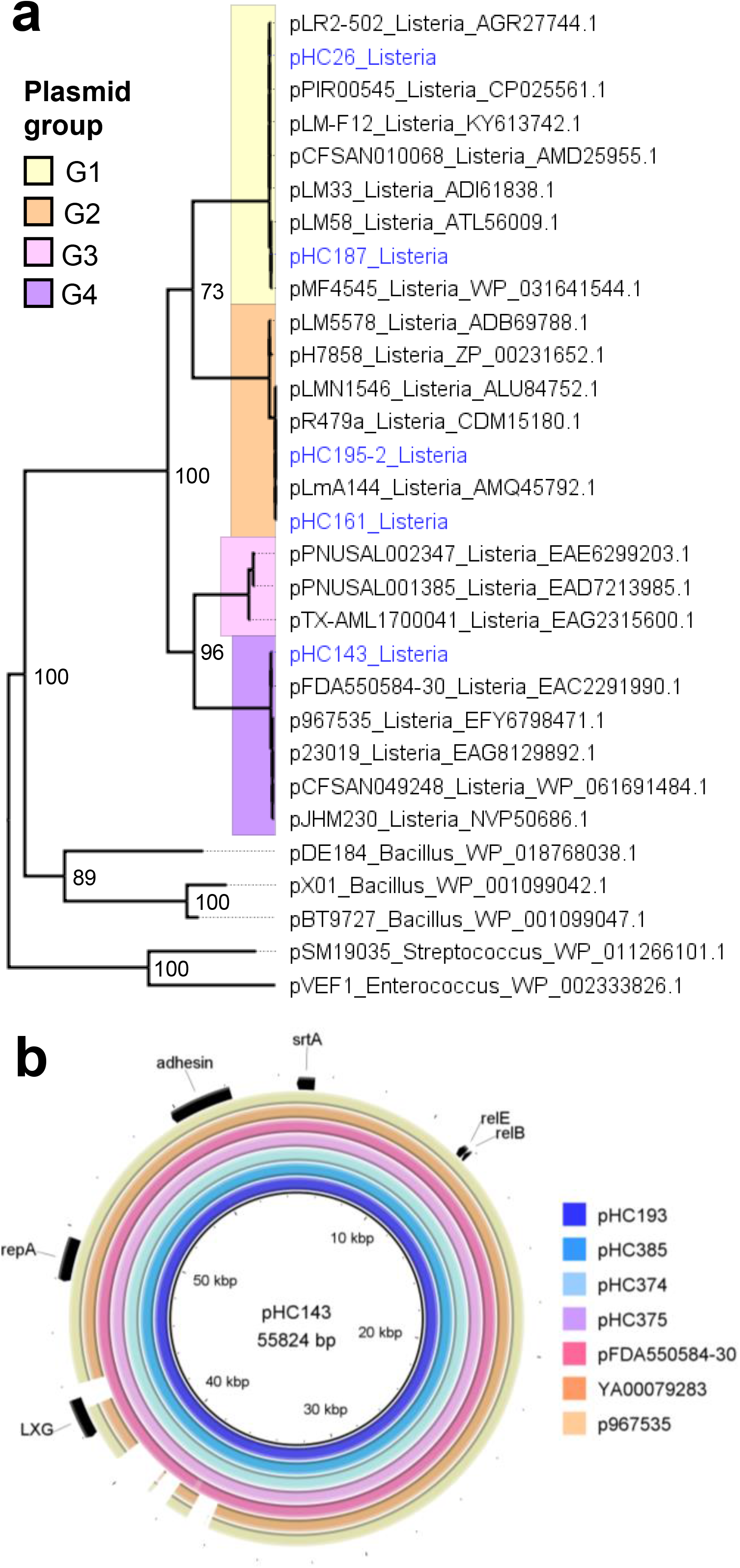
Characterization of plasmids based on RepA. **a** Maximum-Likelihood phylogenetic analysis of the >50 Kb plasmids detected in the present study, based on the *repA* amino acid sequences. The analysis employed the Jones-Taylor-Thornton substitution model with 100 bootstraps and was performed using the MEGA7 software. Bootstrap support values above 70 are shown. Plasmids represented three phylogenetic clades (plasmid groups G1, G2 and G4). Plasmid groups correspond to the groups established by Kuenne et al., (2010) and Schmitz-Esser et al., (2021). Tip labels correspond to plasmid names and host genera; plasmids from this study are labelled in blue. **b** G4 plasmids of the *L. monocytogenes* strains HC193 (this study), HC374 (this study) and FDA550584-30 (SAMN02923676) aligned with >95% identity across the entire length of pHC143 from this study; plasmids of the *L. monocytogenes* strains 967535 (SAMN15680309) and YA00079283 (SAMN08970420) aligned with >95% identity to most of pHC143. The alignment was generated using the BRIG 0.95. For pHC143, plasmid length in base pairs (bp) is given.

Plasmid groups G1 and G2 composed of several well-characterized *L. monocytogenes* reference plasmids that are common in food processing environments (Kuenne et al., 2010; Hingston et al., 2019; Nadiz et al., 2019). pHC143 represented G4, a novel group of *Listeria* plasmids (Schmitz-Esser et al., 2021). The pHC143 plasmid was detected in five isolates of this study, belonging to STs 6 and 149 (see Supplementary Data S1). These STs are hypervirulent, based on the presence of pathogenicity islands LIPI-3 (ST6) and LIPI-4 (ST149) (Fig. 2). Visualisation of assembly graphs indicated that pHC143 was successfully assembled into a single 55.8 Kb contig in all five isolates. pHC143 contained no biocide or heavy metal resistance genes. However, we identified three variants of pHC143 among short-read sequence assembles deposited in GenBank, all of which contain resistance genes against biocides (Fig. 4b, see Supplementary Fig. S1). The first variant contains a benzalkonium chloride resistance cassette (*bcrABC*) and a mercuric resistance (*mer*) operon. The second variant contains a multidrug exporter putatively conferring resistance against quaternary ammonium compounds (*qacC*/*qacH;* WP_000121134.1). The third variant contains the *qacC*/*qacH* gene and a Tn*554*-family transposon carrying an arsenic resistance operon (*arsABCD*). This Tn*554*-family transposon was identified previously in the chromosomes of *L. monocytogenes* (Kuenne et al., 2013). All G4 plasmids contained a predicted fimbrial adhesin (WP_061691480.1), suggestive of a role associated with attachment and host colonisation (Ageorges et al., 2020).

Assembly graphs of pHC192 suggested that the plasmid was also closed successfully into a single 4.6 Kb contig. pHC192 did not contain replication proteins related to the RepA of *Listeria* plasmid groups G1-G4, so the phylogeny of this plasmid was analysed using RepB (Fig. 5a). Phylogenetically, pHC192 clustered closely with plasmids from *Lactobacillus*. Indeed, RepB of pHC192 (WP_035147907.1) was also detected in *Lactobacilli* and *Brochothrix* (100% amino acid sequence identity), suggestive of a broad host range for this plasmid. The closest relative of pHC192 in *Listeria* was the plasmid of the *L. monocytogenes* strain CFIAFB20130002, which possesses the lincosamide resistance gene *lnuA* (WP_001829870.1). Notably, RepB of pHC192 bore no similarity to the replication proteins of the small *Listeria* plasmids pIP823 (WP_172694646.1) and pDB2011 (WP_020277964.1) and shared only 45% amino acid identity with the RepB of pLMST6 (WP_061092472.1). Like pHC192, pLMST6 appears to also have a broad host range, as 100% identical homologues of pLMST RepB (WP_061092472.1) were detected in *Listeria, Salmonella* and *Enterococcus*. These findings suggest that pHC192 and pCFIAFB20130002 represent a novel phylogenetic group of small *Listeria* plasmids (G5) that are distinct from the pLMST6 family of plasmids (G6), implying that several phylogenetically unrelated small plasmids have been acquired by *Listeria* through distinct transfer events across host species.

**Fig. 5.**
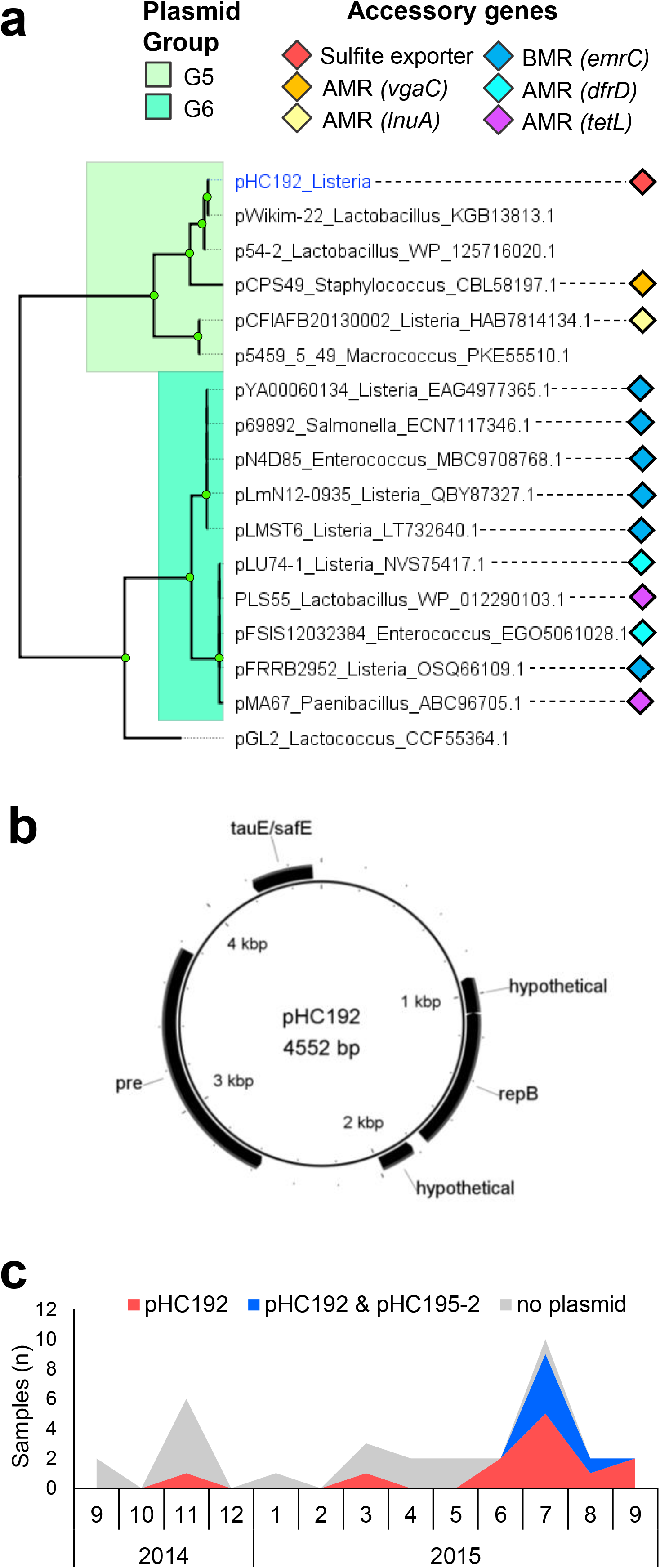
Phylogeny, gene content and epidemiology of the novel plasmid pHC192. **a** Maximum-Likelihood phylogenetic analysis of the <10 Kb plasmids detected in the present study and other related plasmids the *repB* amino acid sequences. Plasmids other than pHC192 were identified and obtained from GenBank using BLASTp. The analysis employed the Jones-Taylor-Thornton substitution model with 100 bootstraps and was performed using the MEGA7 software. Node labels indicate bootstrap support values above 70. Tip labels correspond to plasmid names and host genera; plasmids from this study are labelled in blue. Tip shapes depict harbourage of resistance genes against antimicrobials (AMR) and biocides (BCR). **b** The 4.5 Kbp plasmid pHC192, carrying a putative SafE/TauE family sulphite exporter (WP_016896343.1). The figure was constructed using the BRIG 0.95. Plasmid length in base pairs (bp) is given. **c** Number of samples containing no plasmid, pHC192, or both pHC192 and pHC195-2 among persistent clade C7 during each month of sampling. Plasmid prevalence in C7 increased over the one-year sampling period.

The plasmid pHC192 contains a putative *tauE/safE* -family sulphite exporter gene (WP_016896343.1) (Fig.5b) that is not typically present in *Listeria* plasmids (Hingston et al., 2019; Schmitz-Esser et al., 2021). The sequencing depth of coverage for pHC192 was approximately five times that of the chromosome, suggesting that pHC192 is a high copy number plasmid. This plasmid became increasingly prevalent among persistent clade C7 isolates during the sampling period and was detected in all isolates at the end of the study (Fig. 5c). An additional plasmid, pHC195-2, was detected in several isolates in the latter part of the study period. The pHC195-2 plasmid belonged to the phylogenetic group G2 (Fig. 4a) and closely resembled the reference plasmid pLMR479a (see Supplementary Fig. S2). The acquisition of these plasmids during the course of persistence suggests that they play a role in the adaption of the pathogen to the farm ecosystem.

### Dairy farm isolates of *L. monocytogenes* share common integrative mobile elements with *Enterococci*

Among the 250 dairy farm isolates we identified six chromosomally located mobile elements: the *L. monocytogenes* IS3-like elements (Kuenne et al., 2013); *Listeria* Genomic Island 2 (LGI-2) (Lee et al., 2017); Tn*5801*_B23 (León-Sampedro et al., 2016); and three novel mobile elements, which were submitted to the Transposon Registry (Tansirichaiya et al., 2019), and assigned the labels Tn*7101*, Tn*7103* and Tn*7104*. The elements ICELm1 (Kuenne et al., 2013), LGI-1 (Gilmour et al., 2010), LGI-3 (Palma et al., 2020), Tn*5422* (Lebrun et al., 1994), Tn*554* (Kuenne et al., 2013), Tn*6188* (Müller et al., 2013), Tn*6198* (Bertsch et al., 2013), and chromosomally located Tn*5422* (Lebrun et al., 1994) were not detected.

The IS3-like transposon was significantly more prevalent among isolates in persistent than nonpersistent clades (Fig. 3b). The IS3-like transposon consists of two insertion sequences in Lineage I (IS-1 and IS-2) and a single insertion sequence in Lineage II (IS-3) (Fig. 2). These elements harbour multiple surface-associated lipoproteins, which may facilitate attachment and invasion (Kuenne et al., 2013). The suggested role of the IS3-like transposon in *L. monocytogenes* virulence remains to be determined.

The integrative and conjugative elements (ICEs) LGI-2 and Tn*5801*_B23 were significantly more prevalent among persistent clade isolates than singleton isolates (Fisher’s Exact Test, *p*<.01) (Fig. 3b). LGI-2 carries cadmium and arsenic resistance cassettes and two multidrug transporters (see Supplementary Fig. S3). Identical (100% nucleotide identity) LGI-2 were present among all ST14 and ST145 isolates of this study (Fig. 2). Moreover, BLASTn search identified identical LGI-2 in 11 *L. monocytogenes* and two *Enterococcus faecalis* complete genomes (see Supplementary Table S2), suggestive of recent transfer between these species.

Tn*5801*_B23 was detected in a subset of ST20 isolates, including the persistent clades C9 – C12 (Fig. 2, see Supplementary Data S1). Tn*5801*_B23 detected in this study shared 97% identity with the Tn*5801*_B23 of the *E. faecalis* strain JH2-2 (see Supplementary Fig. S4). Tn*5801*_B23 contains putative resistance genes against the antimicrobial bacitracin (*bcrABD*) and a two-component system (*baeSR*) potentially involved in the regulation of the *bcrABD* operon (León-Sampedro et al., 2016). Unlike Tn*5801*_B23, other Tn*5801*-like elements mediate tetracycline resistance in *Enterococcus, Listeria* and several other Firmicute species (León-Sampedro et al., 2016). In *L. monocytogenes* ST20, Tn*5801*_B23 was inserted downstream of *guaA* (*lmo1096*), which is also the insertion site of the related element ICELm1 of the *L. monocytogenes* strain EGD-e, harbouring cadmium resistance genes (Kuenne et al., 2013).

The putative integrative and mobilizable element (IME) Tn*7101* was detected in the ST155 singleton isolate HC258, where it was inserted between homologues of *lmo2596* and *lmo2597* (see Supplementary Fig. S3). Tn*7101* contains resistance genes against cadmium (*cadA*, *cadC*) and an arsenate reductase (*arsC*). Through a BLAST search we identified a variant of the Tn*7101* containing a seven gene arsenic resistance cassette. This variant, labelled Tn*7102*, was detected in several *L. monocytogenes* and *Enterococcus* genomes deposited in GenBank (see Supplementary Fig. S2). The Tn*7101* and Tn*7102* of *Listeria* and *Enterococcus* were identical (100% nucleotide identity), suggestive of recent promiscuity between the two genera. Arsenic resistance genes in Tn*7102* were distantly related (≥67% identity) to the arsenic resistance cassette of LGI-2 (see Supplementary Fig. S3).

The putative IME Tn*7103* was detected in the ST119 singleton isolate HC183, where it was inserted between *lmo0810* and *lmo0811*. This transposon contained putative virulence genes encoding an InlJ-like internalin and a bacterial immunoglobulin (Big)-like protein (see Supplementary Fig. S5). A BLAST search confirmed the presence of Tn*7103* in other *L. monocytogenes* strains, including N12-2532 (SAMN09947958), but we did not identify this element in other species.

The putative ICE Tn*7104* was detected in the ST391 singleton isolate HC187 and was inserted between *lmo1786* and *lmo1787.* This transposon contained a putative type I restriction-modification system (see Supplementary Fig. S6). Tn*7104* was identified in several other *L. monocytogenes* strains deposited in GenBank, including the *L. monocytogenes* ST391 strain SHL013 (SAMN03265960), but we did not identify this element in other species.

### A novel prophage harbouring cadmium resistance genes was identified in a persistent clade of *L. monocytogenes*

All 250 dairy farm isolates from this study contained the *L. monocytogenes* monocin (Zink et al., 1995), and 0 – 3 additional prophages, which were detected at eight insertion sites (Fig. 3c). Prophages inserted into tRNA-Arg(tct) were significantly more prevalent among isolates in persistent clades, and prophages inserted into tRNA-Lys(ctt) were significantly more prevalent among nonpersistent clades (Fisher’s Exact Test, *p*<.05).

OPTSIL taxonomic clustering assigned prophages from this study into six genera. Prophages inserted into *comK* and tRNA genes were assigned to genera of *Siphoviridae* that are known to only infect *Listeria*. Surprisingly, in the isolate HC189, a 67 Kb *Myovirus* was inserted into *comK*, a site usually occupied by *Siphoviridae* (Pasechnek et al., 2020).

Prophages inserted between the *rlmCD* (*lmo1703*) and *fosX* (*lmo1702*) genes were not related to any of the *Listeria* specific phage genera, but instead represented a separate genus that infects several Firmicute species (Fig. 6). Many of the phages in this genus harbour antimicrobial and heavy metal resistance cassettes (see Supplementary Fig. S7). In this study, phages inserted between the *rlmCD* and *fosX* were detected among all isolates of persistent clade C8 and among three singleton isolates (see Supplementary Data S1). Among isolates of persistent clade C8, prophages inserted between *rlmCD* and *fosX* all harboured a cadmium resistance cassette (see Supplementary Fig. S7). In contrast, in the singleton isolates prophages inserted between *rlmCD* and *fosX* harboured no cadmium or antimicrobial resistance genes. Within *Listeria* genomes deposited in GenBank, we identified prophages inserted between *rlmCD* and *fosX* that carried resistance genes against cadmium (*cadA*), macrolides (*mefA*, *msrD*), tetracycline (*tetM*) and streptogramin (*vatA*).

**Fig. 6.**
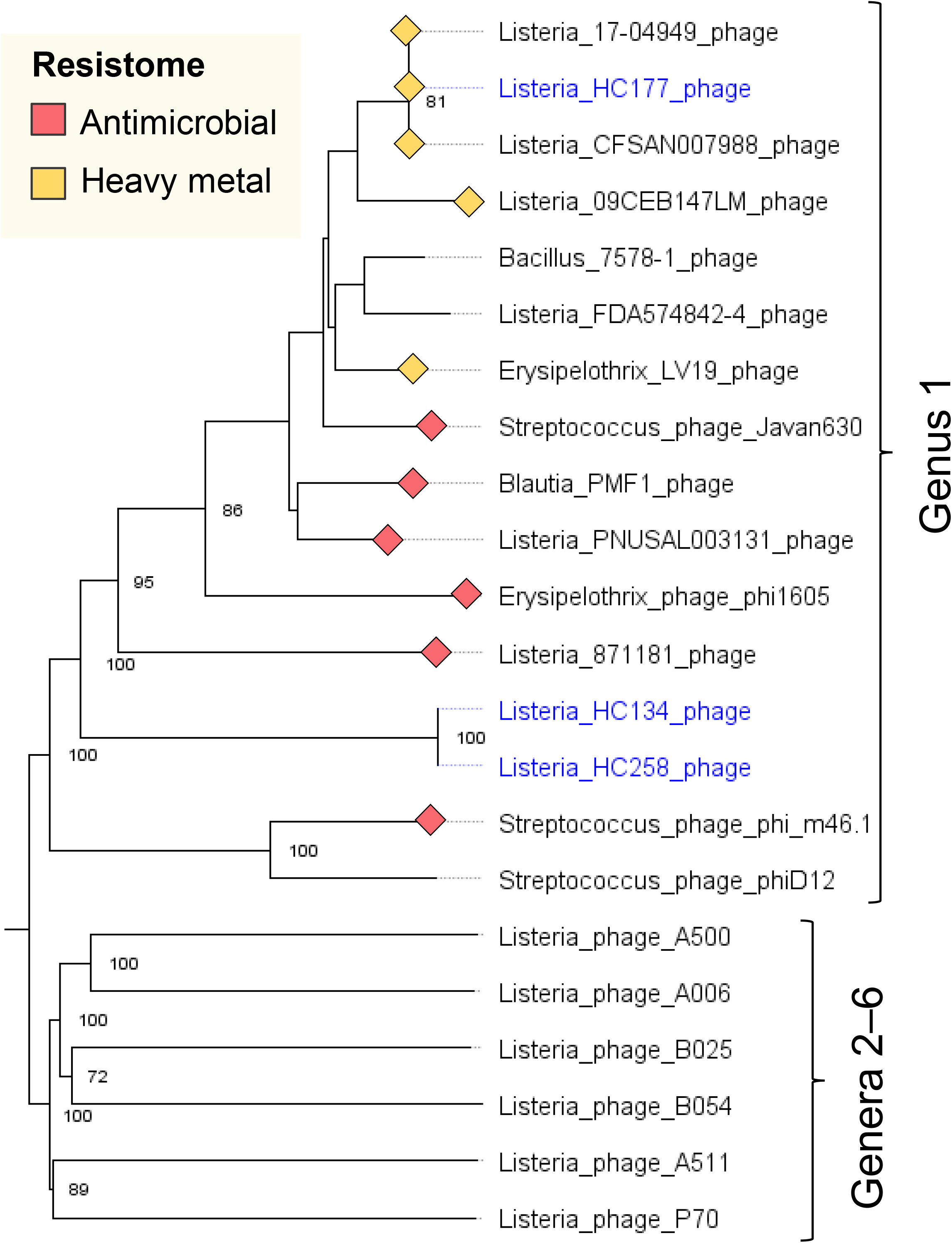
Prophages inserted between *rlmCD* and *fosX* belonged genus of *Siphovirus* having a broad host species range and a tendency to harbour antimicrobial or heavy metal resistance genes. Minimum evolutionary tree and taxonomic clustering of six *Listeria* specific phages (genera 2-6); two prophages from this study that were inserted between *rlmCD* and *fosX* genes (genus 1, blue); and prophages from *Listeria* and other Firmicute species obtained from GenBank (genus 1, black). Phylogenetic analyses and clustering were generated with the Victor online tool (https://victor.dsmz.de), using the model D6 and 100 bootstrap replicates. The tree was visualized using FigTree 1.4.4. Bootstrap support values above 70 are shown.

### Systems that protect against invading DNA were negatively associated with the persistence of *L. monocytogenes* on dairy farms

A genome wide association study was conducted to assed which genes were associated with persistent versus nonpersistent clades. Because no persistent clades belonged to Lineage I, the analysis was restricted to Lineage II. Among the genes that were positively associated with persistence on dairy farms were a gene associated with biofilm formation (*bapL*), a peptidoglycan hydrolase (*murA*) (see Supplementary Table S3). Interestingly, *bapL* has previously been implicated in the adaptation of *L. monocytogenes* to the food processing environment (Maury et al., 2019). In contrast, genes associated with the CRISPR-*cas* type IIA system, and the type II restriction-modification system LmoJ3 (Lee et al., 2012) were negatively associated with persistence in the dairy farm environment (see Supplementary Table S3). CRISPR-*cas* systems and restriction-modification systems may act in synchrony to protect the host against invading prophages and other mobile elements (Price et al., 2016). Additionally, a putative recombination and DNA strand exchange inhibitor protein (WP_03166494.1) was negatively associated with persistence. These findings agree with the lower prevalence of mobile genetic elements and prophages among isolates in nonpersistent than persistent clades and suggest that systems involved with inhibiting invading DNA are detrimental for the persistence of *L. monocytogenes* in the dairy farm environment.

*L. monocytogenes* hypervariable hotspots 1 and 8 (Kuenne et al., 2013) contained genes both positively and negatively associated with persistence. Hypervariable hotspot 1 consists of a ESX-1 like type VII secretion system (T7SS) that has a suggested role in bacterial antagonism (Bowran & Palmer, 2021). Indeed, most genes associated with persistence belonged to *L. monocytogenes* hypervariable hotspots or prophages, suggesting that the role these components play in *Listeria* niche adaptation deems further study.

## Discussion

Whole genome sequencing and subsequent analyses of 250 *L. monocytogenes* isolates from dairy farms illustrated that dairy farm isolates are hosts to a diversity of mobile genetic elements that carry, or have the potential to carry, resistance genes against antimicrobials, biocides and heavy metals. We found that prophages and other mobile genomic elements were significantly more numerous among isolates belonging to persistent than nonpersistent clades. Moreover, systems that provide immunity against invading mobile genetic elements (Garneau et al, 2010; Lee et al., 2012; Price et al., 2016), namely the CRISPR-*cas* IIA system, the type II restriction-modification system LmoJ3, and a putative recombination and DNA strand exchange inhibitor protein, were negatively associated with persistence. These findings suggest that mobile elements may support the persistence of *L. monocytogenes* inhabiting farms. Many of the mobile elements we identified carried genes encoding phenotypes that promote the survival of *L. monocytogenes* on farms, such as antimicrobial resistance genes or virulence factors. Moreover, genes responsible for the conjugation of mobile elements may have a dual role in promoting biofilm formation and invasion of the mammalian host (Barrios et al., 2005; Ageorges et al., 2020), thereby supporting the survival of *L. monocytogenes* on farms.

We identified a surprising diversity of mobile genetic elements encoding heavy metal resistance genes among the dairy farm isolates and acquired heavy metal resistance genes were more common among isolates in persistent than nonpersistent clades. Similarly, heavy metal resistance genes are more prevalent among persistent than nonpersistent *L. monocytogenes* subtypes from foods and food processing environments (Harvey & Gilmour, 2001; Pasquali et al., 2018). Whether heavy metal resistance genes contribute to persistence or merely co-occur with other fitness enhancing determinants remains unclear (Parsons et al., 2020). Nevertheless, heavy metal resistance genes may represent useful markers to aid the detection of *L. monocytogenes* strains with fitness-enhancing mobile genetic elements.

In the present study we found a novel plasmid (pHC143; plasmid group G4) that infected hypervirulent subtypes of *L. monocytogenes*. Although pHC143 was devoid of biocide and heavy metal resistance genes, such genes are common on other G4 plasmids infecting hypervirulent ST1 and ST6 strains (Schmitz-Esser et al., 2021). Indeed, we noted that a G3 plasmid harbouring the biocide resistance gene *qacC* and the arsenic resistance cassette *arsABCD* in the ST6 outbreak isolate YA00079283, associated with the largest listeriosis outbreak known to date (Smith et al., 2019).

We identified four transposons in *Listeria*, namely LGI-2, Tn*5801*_B23, Tn*7101* and Tn*7102*, that closely resembled transposons in *Enterococci*, suggestive of recent transfer between the two genera. The co-occurrence of genomic elements in *Enterococci* to *Listeria* was unsurprising, as both genera are highly prevalent in animal faeces and farms (Franz et al., 1999; Hellström et al., 2010; Castro et al., 2018). Transfer of conjugative elements has been demonstrated both from *Enterococci* to *Listeria* and vice versa (Jahan & Holley, 2016; Haubert et al., 2018) indicating that both genera are potential donors. The extent to which *Enterococci* and other Firmicutes contribute to the horizontal spread of mobile elements and their associated antimicrobial, biocide and heavy metal resistance determinants in *Listeria* has implications for food safety and should be explored through further study.

Bacitracin resistance genes, mediated by Tn*5801*_B23, were common among *L. monocytogenes* from all three farms investigated. Moreover, Tn*5801*_B23 was significantly more prevalent among isolates in persistent clades than nonpersistent clades. The widespread use of bacitracin as a growth promoter in animal feeds has facilitated the expansion of bacitracin resistance in *Enterococci* (Aarestrup et al, 2000; Chen et al., 2016), and probably also *L. monocytogenes*, as animal feeds are frequently contaminated by *Listeria* (Hellström et al., 2010; Castro et al., 2018). Nevertheless, the frequent detection of Tn*5801*_B23 in this study remains curious, as feed supplementation with bacitracin subsided in Finland in the 1990’s (Aarestrup et al., 2000).

We found that prophages were more prevalent among persistent clade isolates than singleton isolates. It is unclear whether the higher number of prophages among persistent clade isolates represents a beneficial role for these elements or is a side effect of being receptive to foreign DNA. There is increasing evidence that prophages can mediate beneficial phenotypes for their host. Phages mediate resistance or virulence properties in numerous bacterial species (Harrison & Brockhurst, 2017), and in *Listeria*, *Siphoviruses* inserted into *comK* were found to regulate the gene in a symbiotic manner (Pasechnek et al. 2020). Here, we discovered phage-mediated carriage of cadmium resistance and various antimicrobials in *Listeria*, suggesting that prophages contribute to the spread of phenotypes supporting persistence. Moreover, we noted that these phages belonged to a genus of *Siphovirus* with an apparently broad host species range that were introduced to *Listeria* through several distinct transfer evets. Host species jumps have the potential accelerate the transfer of novel resistance determinants between *Listeria* and other Firmicutes.

It is worth noting that not all persistent clades harboured mobile elements, suggesting that other factors also contribute to the survival of *L. monocytogenes* on dairy farms. We found that genes putatively involved in biofilm formation (*bapL*) and interbacterial competition (T7SS), which are not located in mobile elements, were significantly associated with persistence. In addition, the predominance of persistent *L. monocytogenes* strains in the dairy farm environment is associated with inadequacies in production hygiene (Castro et al., 2018). Therefore, the persistence of *L. monocytogenes* in the dairy farm environment is likely the result of a multifactorial combination of bacterial and environmental factors.

In conclusion, our study indicates that *L. monocytogenes* inhabiting the dairy farm environment are receptive to a diversity of prophages and mobile genetic elements. We suggest that mobile elements enable *L. monocytogenes* to adapt to the stresses encountered in the farm ecosystem and in general improve the fitness of the pathogen on farms, thereby supporting persistence. Given the abundance of *L. monocytogenes* on farms (Nightingale et al., 2004; Esteban et al., 2009; Castro et al., 2018) and the apparent exchange of mobile genetic elements between *Listeria* and other Firmicute species, *L. monocytogenes* occurring in agroecosystems should be viewed as a potential reservoir of mobile genetic elements. Importantly, many of these elements have the potential to carry and spread antimicrobial, biocide, and heavy metal resistance genes. The spread of mobile genetic elements and resistance determinants from primary production to *Listeria* in the food processing environments has important food safety implications and should be explored further. The present study represents a step forward in this effort and in our understanding of listerial ecology in the agroecosystem.

## Methods

### Whole genome sequencing

Altogether 250 *L. monocytogenes* isolates obtained from three Finnish dairy cattle farms during 2013–2016 (Castro et al., 2018) were selected for whole genome sequencing in the present study (see Supplementary Data S1). DNA was extracted from overnight cultures using the guanidium thiocyanate extraction method (Pitcher et al. 1989). DNA samples were standardized to a concentration of 10 ng/μl using the dsDNA BR Assay Kit (Thermo Fisher Scientific; Waltham, MA, USA) using the Qubit Fluorometer (Thermo Fisher Scientific). Genomic libraries were constructed form the DNA samples using the Nextera XT DNA Sample Preparation Kit (Illumina; San Diego, CA, USA), and paired-end sequencing (2×250 bp) was performed using the Illumina HiSeq platform.

### Genome assembly, pangenome construction, subtyping

Following the removal of adapter sequences and low-quality reads using Trimmomatic 0.36 (Bolger et al., 2014), draft genomes were assembled using SPAdes 3.9 with K-mer values 55, 77, 99, 113 and 127 (Bankevich et al., 2012). Assembly quality was assessed using QUAST 4.0 (Gurevich et al., 2013) and taxonomic assignment was performed using Kraken (Wood & Salzberg, 2014). The assemblies were annotated using Prokka 1.12 (Seemann, 2014). The pangenome of the sequenced isolates was constructed using Roary 3.8.0 (Page et al., 2015) with the protein identity cut-off value set at 90%. Multi Locus Sequence Types (ST) were determined *in silico* from the assembled genomes using the BIGSdb-*Lm* database (Moura et al., 2017), which utilizes the schema developed by Ragon et al. (2008). The BIGSdb-*Lm* database was also used to identify pathogenicity islands associated with hypervirulence (LIPI-3, LIPI-4), and genes associated with antimicrobial and biocide resistance among the assembled genomes. Genome assemblies were deposited in GenBank under the BioProject number PRJNA704814 (see Supplementary Data S1).

### Maximum-Likelihood Phylogenomic Analysis

Phylogenomic reconstruction of the 250 *L. monocytogenes* isolates was performed using the Lyve-SET 1.1.4f pipeline (Katz et al., 2017), using *L. monocytogenes* EGD-e genome (NC_003210.1) as reference. The Lyve-SET pipeline was run using *Listeria monocytogenes* pre-sets (Katz et al., 2017), with additional options --mask-phages, --mask-cliffs, and -- read_cleaner CGP. In brief, the pipeline generated genome alignments by mapping quality-filtered reads to a reference genome. To improve the accuracy of phylogenomic inference, putative prophage genes were removed from the reference genome prior to mapping. Mapping was followed by the detection of high-quality SNPs, having ≥ 10x depth of coverage and ≥ 75% consensus among reads. Recombinant sites within the genome alignments generated by Lyve-SET were identified and removed using Gubbins 3.0 (Croucher et al., 2015). PhyML 3.3 (Guindon et al., 2010) was used to infer Maximum-Likelihood -phylogeny of each ST using a general time reversible model (GTR) with 100 bootstrap replicates.

In addition, the phylogeny of each ST harbouring putative persistent clades was inferred individually. Persistent clades of *L. monocytogenes* were defined monophyletic clades of isolates with PWDs < 20 SNPs (Pightling et al., 2018) that were isolated from the same farm from ≥3 samples during ≥6 months. For each ST, a draft assembly from the present study with the best quality statistics, i.e. the highest N50 value and lowest number of contigs (see Supplementary Data S1), was used as a reference genome. The phylogenomic analyses were executed as described above using the Lyve-SET pipeline, Gubbins and PhyML.

### Detection and analysis of plasmids

Plasmids were identified by aligning the whole genome assemblies against *Listeria* plasmids deposited in GenBank with the aid of BLASTn (http://www.ncbi.nlm.nih.gov/blast). Alignments were inspected manually. Additionally, whole-genome assembly graphs generated by SPAdes were visualized using Bandage 0.8.1 (Wick et al., 2015) and extrachromosomal elements were inspected manually. Maximum-Likelihood phylogeny of the plasmids, based on the amino acid sequence alignments of the *repA* gene, were generated with MEGA7 (Kumar et al., 2016), using the Jones-Taylor-Thornton substitution model with 100 bootstraps. Alignments of the amino acid sequences of the *repB* gene were used to compare plasmids in which *repA* was absent. Plasmid alignments were generated and visualized using BRIG 0.95 (Alikhan et al., 2018) and EasyFig 1.2 (Sullivan et al., 2011).

### Detection and analysis of chromosomal mobile genetic elements

The occurrence of the chromosomal mobile genetic elements ICELm1 (Kuenne et al., 2013), LGI-1 (Gilmour et al., 2010), LGI-2 (Lee et al., 2017), LGI-3 (Palma et al., 2020), Tn*5422* (Lebrun et al., 1994), Tn*6188* (Müller et al., 2013), Tn*6198* (Bertsch et al., 2013), and the IS3-like and Tn*554*-like transposons of *L. monocytogenes* (Kuenne et al., 2013), among isolates from this study was assed, by aligning the integrases, transposases, recombinases associated with these elements against the pangenome (the pan_genome_reference -file generated by Roary) with the aid of tBLASTtn. Hits were inspected manually. Additionally, the pangenome was searched for annotations including “recombinase”, “integrase”, “transposase”, “transposon”, “cadmium”, “arsenic”, “mercuric”, “*ardA*”, “*ftsK*”, “P60” and “*iap*” and hits were inspected manually. EasyFig 1.2 was used to align and visualize the identified transposons and their occurrence among genomes deposited in GenBank was assessed using BLAST.

### Detection and analysis of prophages

Prophages inserted into the *L. monocytogenes* genomes were identified using PHASTER (Arndt et al., 2016) and the insertion sites were inspected manually. Phylogeny and taxonomic clustering of prophages classified by the PHASTER algorithm as “intact” were inferred using VICTOR (Meier-Kolthoff & Göker, 2017). Nineteen additional *Listeria* phage genomes and one Streptococcal phage genome obtained from GenBank were included in the analyses for reference (see Supplementary Table S1). In brief, VICTOR applies the Genome-BLAST Distance Phylogeny (GBDP) method (Meier-Kolthoff et al., 2013) to obtain pairwise distances, from which balanced minimum evolution trees are inferred. VICTOR utilizes OPTSIL (Göker et al., 2009) to obtain taxonomic clustering. Duplicate phage genomes are removed from the analysis. Trees generated by VICTOR were visualized using FigTree 1.4.4 (http://tree.bio.ed.ac.uk/software/figtree/). BLAST was used to identify phages inserted between *rlmCD* and *fosX* in the genomes of *Listeria* and other bacterial species deposited in GenBank, and hits were inspected manually. Phylogeny and taxonomic clustering of prophages inserted between *rlmCD* and *fosX* were inferred using VICTOR.

### Identification of genes associated with predominance

Scoary 1.6.16 (Brynildsrud et al., 2016) was used to identify genes significantly associated with occurrence in persistent clade isolates versus singleton isolates. Scoary was executed using default options, using the gene_presence_abence.csv -file generated by Roary as input. Associations with a Bonferroni corrected *p*<.05 were considered significant. As all predominant clades belonged to Lineage II, the analysis was limited to the 233 Lineage II isolates of this study to reduce noise arising from population structure bias.

## Supporting information

see Supplementary

## Acknowledgements

This research was supported by the Finnish Ministry of Agriculture and Forestry (grant number 618/03.01.02/2017) and by the Walter Ehrström Foundation. The authors want to acknowledge Esa Penttinen for his assistance with DNA extraction.

## Author Contributions

H.C. conducted the analyses and wrote the manuscript. H.C. and M.L. provided the study materials and designed the study. F.D., H.K. and M.L. supervised the study. All authors participated in data interpretation and reviewed the manuscript.

