## Supplementary material for "Mobile elements habouring heavy metal and bacitracin resistance cassettes are common among *Listeria monocytogenes* persisting on dairy farms": see Supplementary

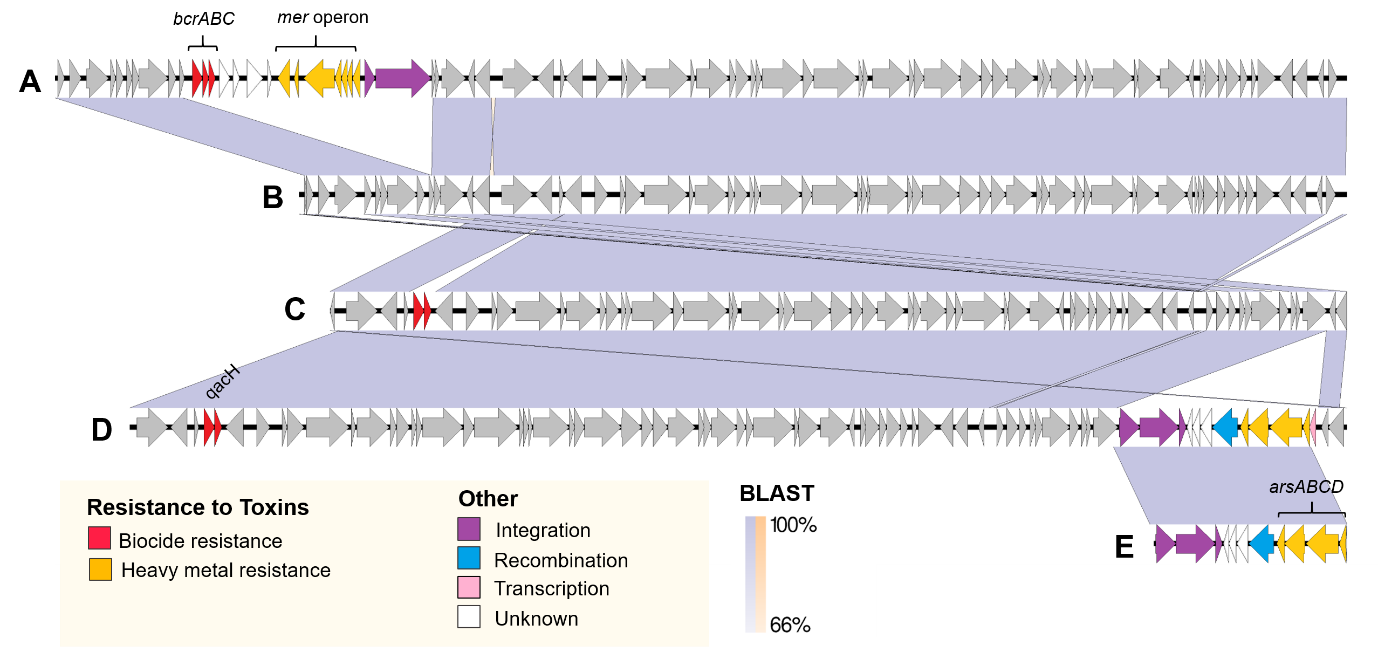

**Supplementary Figure S1. Comparison of group 3 (G3) plasmids**

Alignment of pFDA550584-30 (A), pHC143 (B), p967535 (C), pYA00079283 (D), and the Tn554-like transposon of the strain SLCC2372 (F). pFDA550584-30 contains mercury (*mer* operon) and benzalkonium chloride (*bcrABC*) resistance cassettes; p967535 and pYA00079283 contain the biocide resistance gene *qacH*. Additionally, pYA00079283 contains a transposon carrying the arsenic resistance cassette (*arsABCD*). Colouring reflects functional annotation; genes conserved in all G3 plasmids are coloured in grey.

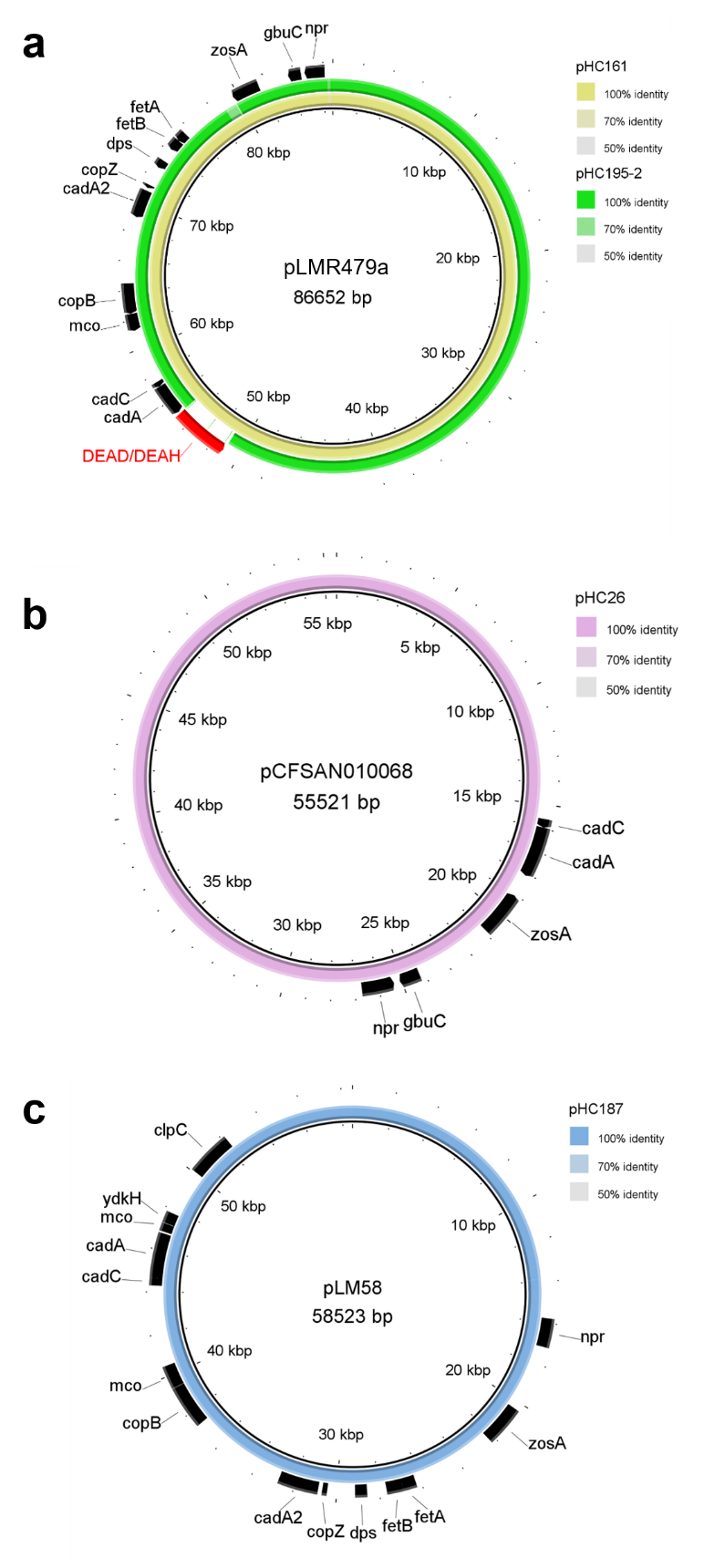

**Supplementary Figure S2. Plasmids of groups G1 and G2 detected in this study**

Plasmids identical to pLMR479a (A), pCFSAN010068 (B), pLM58 (C) were detected in this study. The plasmid pHC195-2 was identical to pLMR479a, except for the absence of a putative DEAD/DEAH box helicase (WP_077913968). Genes putatively associated with heavy metal detoxification or stress tolerance are shown.

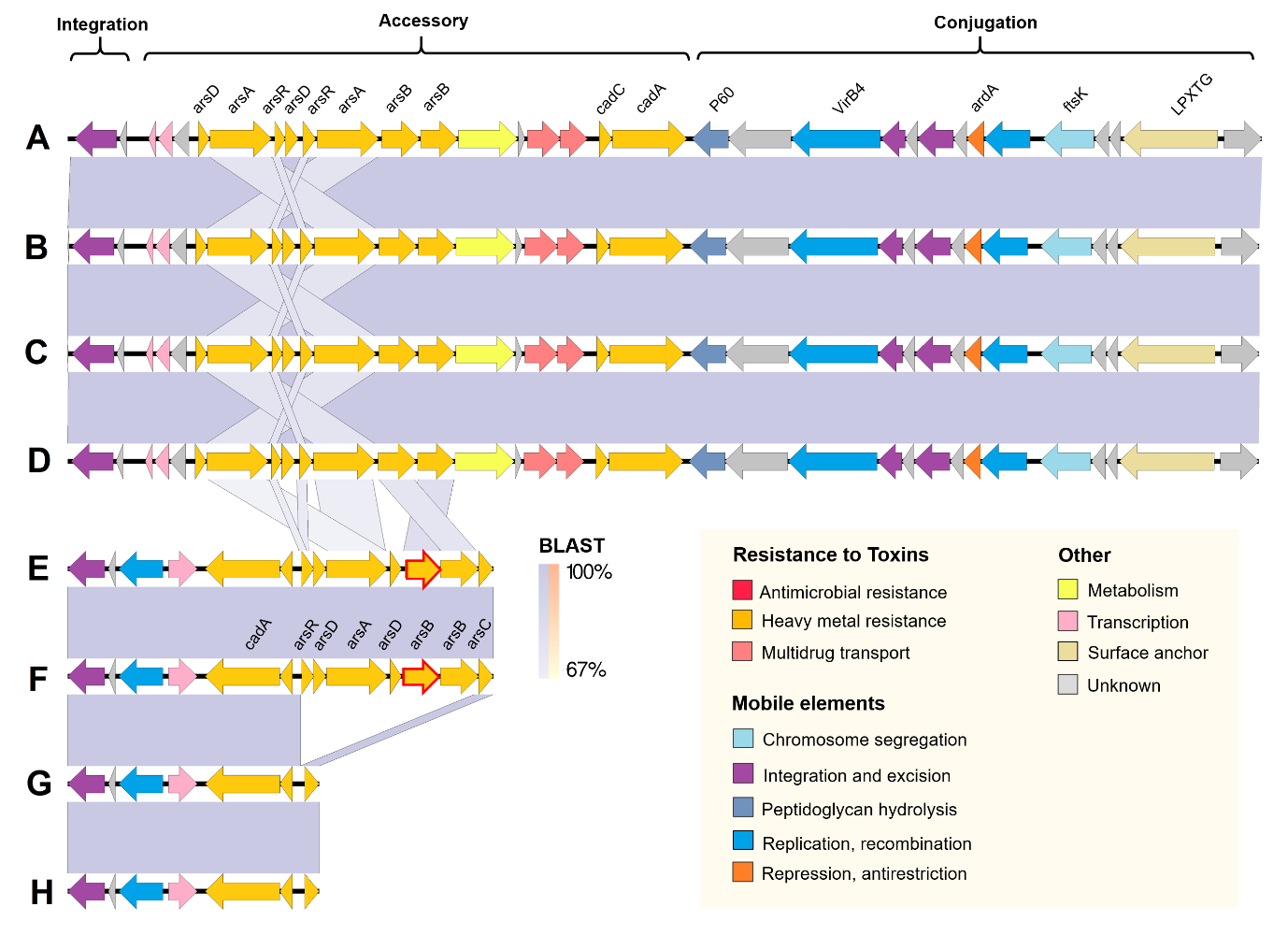

**Supplementary Figure S3. *Listeria* genomic island 2 (LGI-2), Tn*7101* and Tn*7102* were identical in *Listeria* and *Enterococci***

LGI-2 were identical in ST14 (A) and ST145 (B) of this study, in *Listeria* strain J1-220 (C) and in *Enterococcus* strain 110 (D). Tn*7102* were identical in *Listeria* strain ICDC_LM1233 (E) *Enterococcus* strain H112E (F). Tn*7101* were identical in strain HC258 of this study (G) and *Enterococcus* VE80 (H). LGI-2 is an integrative and conjugative element containing an integrase and a type IV secretion system. The novel elements Tn*7101* and Tn*7102* contain ICEBs1_C -like integrases but lack the conjugation infrastructure. Red outlining of arrows indicates pseudogene.

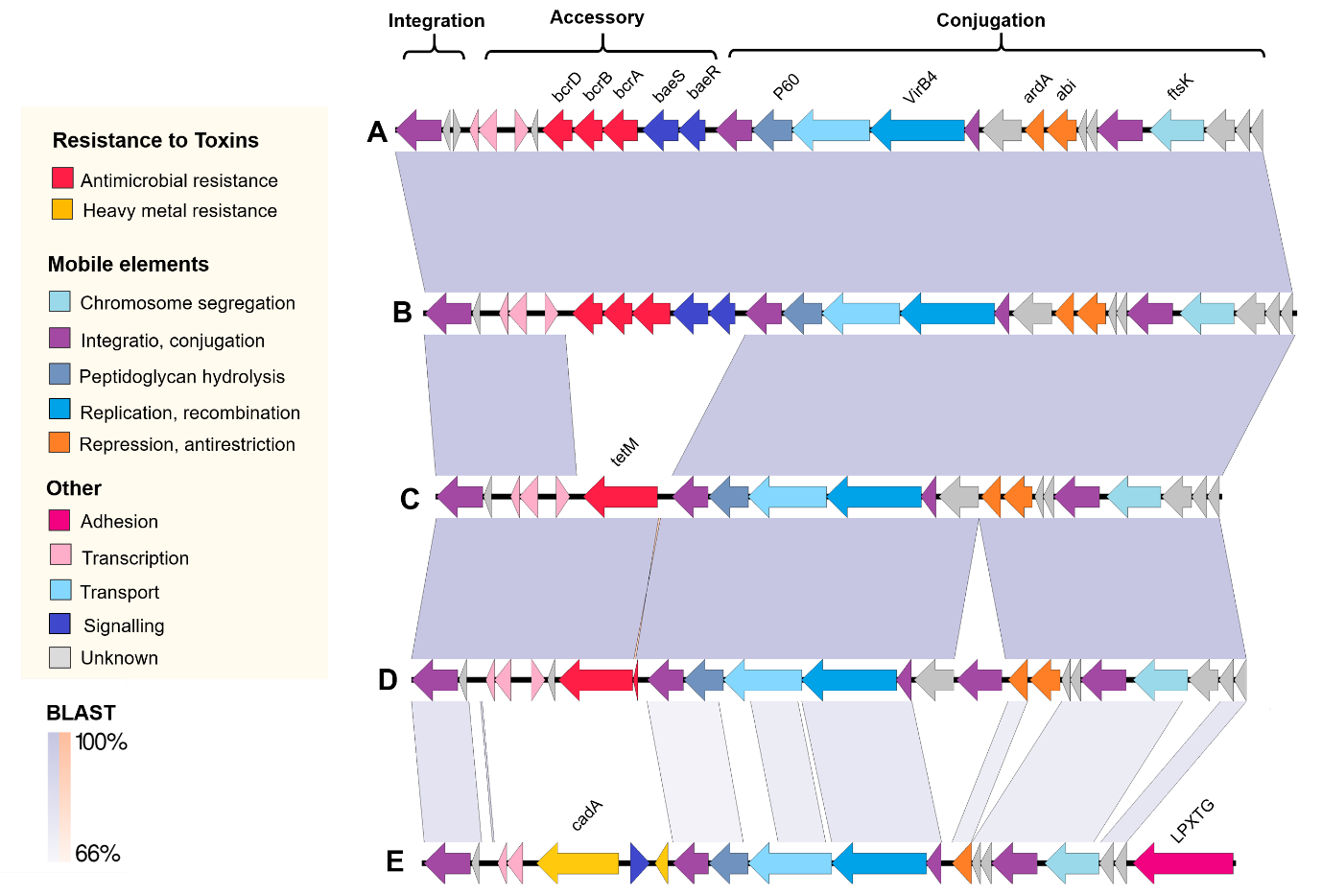

**Supplementary Figure S4. Comparison of Tn*5801*-like integrative and conjugative elements**
Tn*5801*_B23 is identical in *Enterococcus* JH2-2 (A) and the ST20 strains of this study (B). Tn*5801*_B23 contains a bacitracin resistance cassette *bcrABD*. Tn*5801*_B23 is related to Tn*5801*_B15 from *Enterococcus* Ef1 (C); to the Tn*5801*_B15-like transposon of *Listeria* L2624 (D); and to ICELm1 from *Listeria* EGD-e*.*

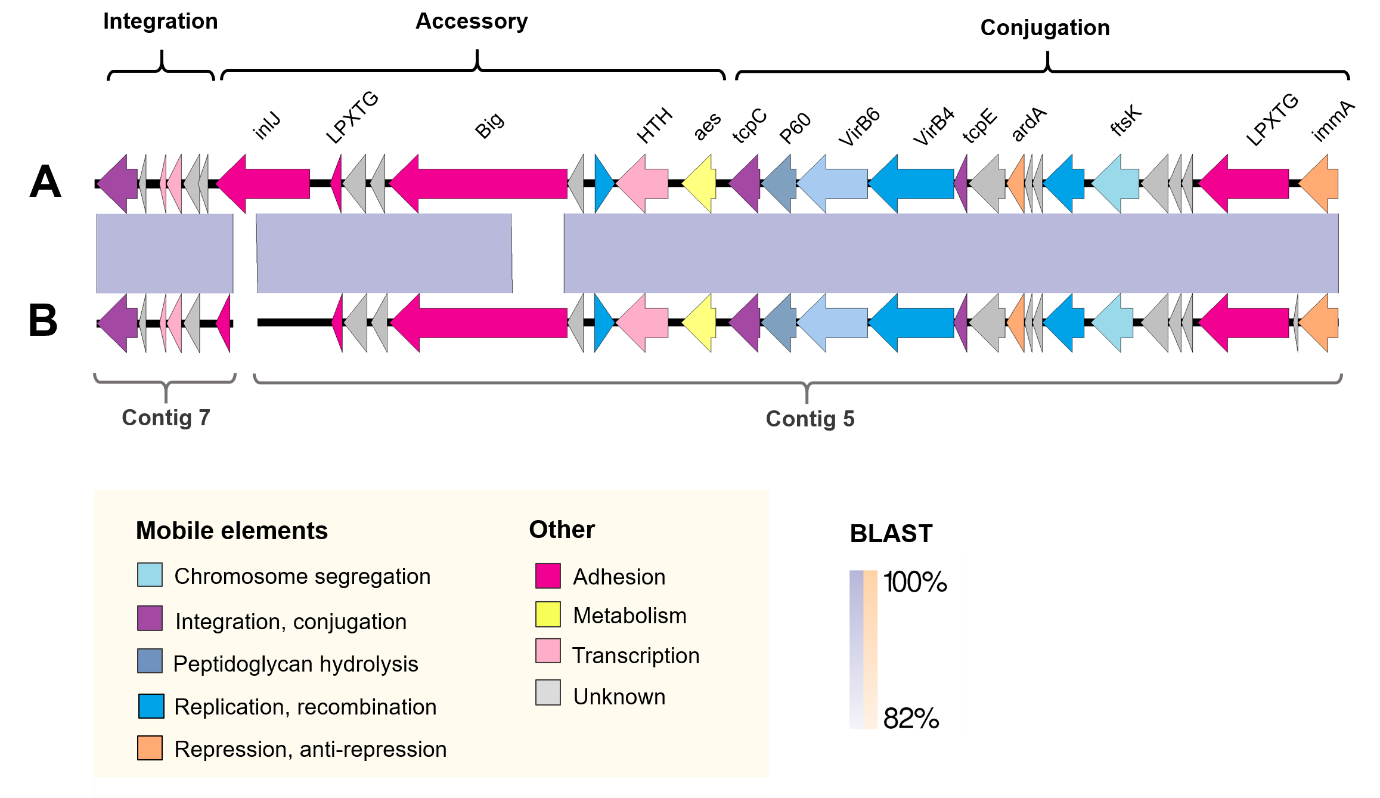

**Supplementary Figure S5. Comparison of Tn*7103***
Tn*7103* of the *L. monocytogenes* strain N12-2532 (A) was identical to HC183 from the present study (B). Tn*7103* is a novel integrative and conjugative element encoding internalin J-like and bacterial immunoglobulin 3-like (Big-3) proteins, putatively associated with attachment and invasion. Red bordering of a gene indicates that it is a pseudogene.

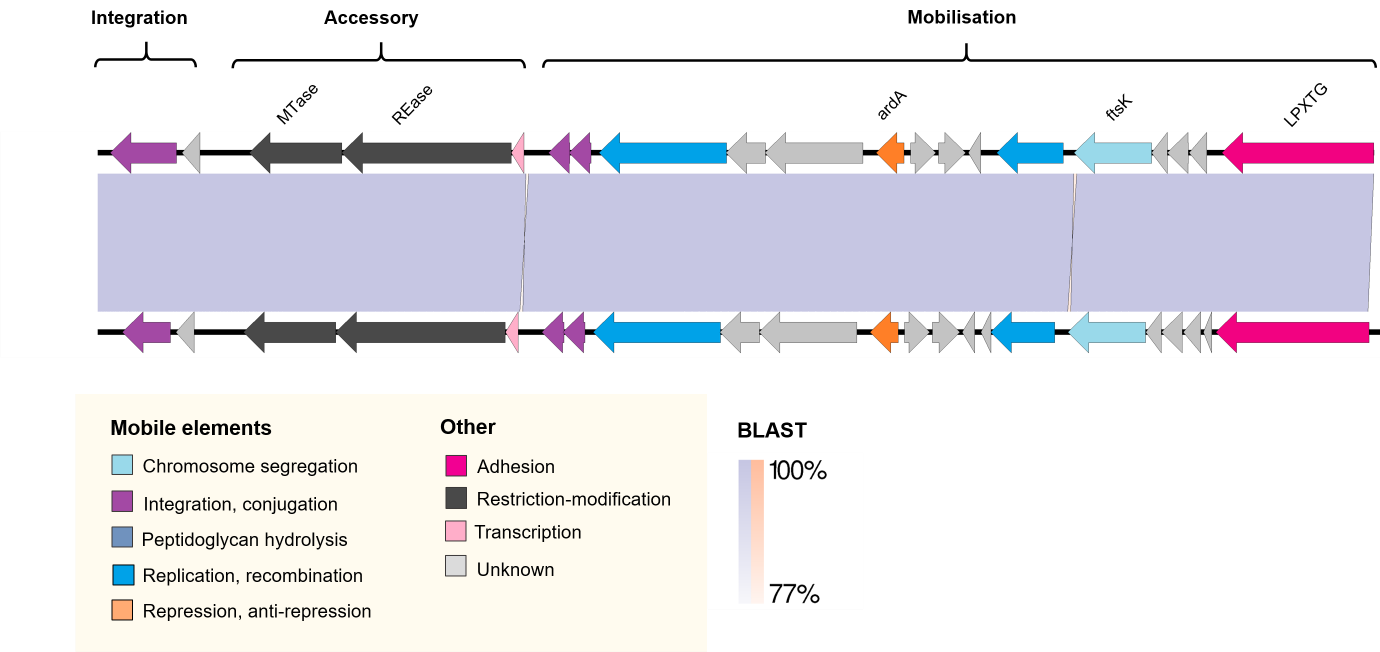

**Supplementary Figure S6. Comparison of Tn*7104***
Tn*7104* of the *L. monocytogenes* ST391 strain SHL013 (A) was identical to that of the ST391 strains of the present study (B). Tn*7104* is a novel mobile element encoding a putative type I restriction-modification system. Tn*7104* contains an integrase but appears to lack a type IV secretion system, suggesting that it is an integrative and mobilizable element (IME). The figure was constructed using EasyFig 1.2.

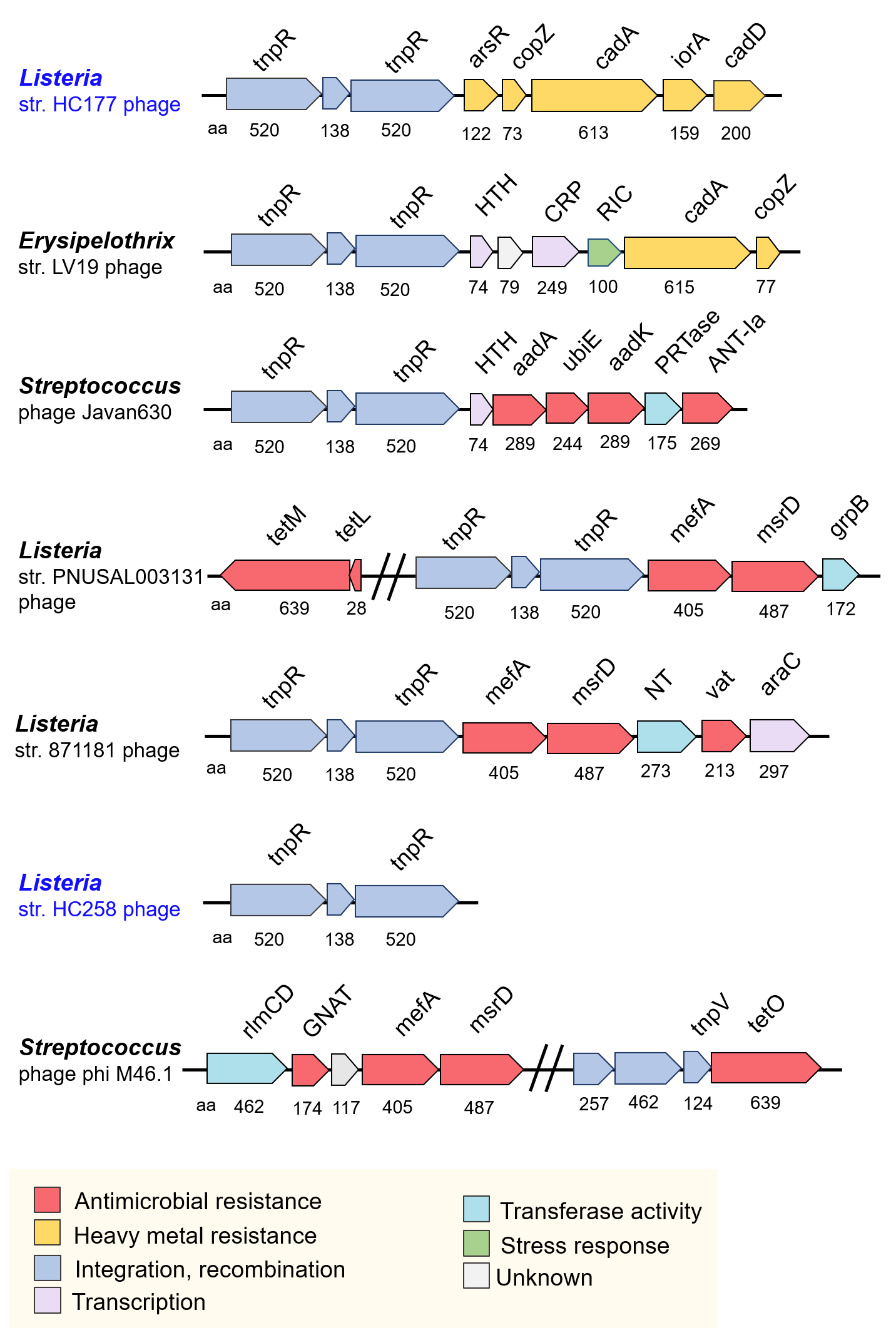

**Supplementary Figure S7. Antimicrobial and heavy metal resistance genes carried by genus 1 prophages**

Schematic representation of the antimicrobial and heavy metal resistance cassettes carried by genus 1 prophages. The amino acid (aa) length is written below each gene; gene name or domain is written on top. Gene colours reflect functional prediction. Isolates form this study are labelled in blue.

**Table S1.** Phage genomes obtained from GenBank that were included the phylogenetic and taxonomic analysis of prophages.

| **Phage name** | **Accession** | **Family** | **Genus** | **Host** |
| --- | --- | --- | --- | --- |
| A006 | NC_009815.1 | *Siphoviridae* | unclassified | *Listeria* |
| A118 | NC_003216.1 | *Siphoviridae* | unclassified | *Listeria* |
| A500 | NC_009810.1 | *Siphoviridae* | unclassified | *Listeria* |
| A511 | NC_009811.2 | *Herelleviridae* | *Pecentumvirus* | *Listeria* |
| B025 | NC_009812.1 | *Siphoviridae* | *Psavirus* | *Listeria* |
| B054 | NC_009813.1 | *Myoviridae* | unclassified | *Listeria* |
| Javan630 | MK448997.1 | *Siphoviridae* | unclassified | *Streptococcus* |
| LP-030-2 | NC_021539.2 | *Siphoviridae* | *Psavirus* | *Listeria* |
| LP-030-3 | NC_024384.1 | *Siphoviridae* | unclassified | *Listeria* |
| LP-037 | NC_021787.2 | *Siphoviridae* | *Homburgvirus* | *Listeria* |
| LP-048 | NC_024359.1 | *Herelleviridae* | *Pecentumvirus* | *Listeria* |
| LP-101 | NC_024387.1 | *Siphoviridae* | *Psavirus* | *Listeria* |
| LP-HM00113468 | MT500540.1 | *Siphoviridae* | *Psavirus* | *Listeria* |
| P100 | NC_007610.1 | *Herelleviridae* | *Pecentumvirus* | *Listeria* |
| P70 | NC_018831.1 | *Siphoviridae* | *Homburgvirus* | *Listeria* |
| PSA | NC_003291 | *Siphoviridae* | *Psavirus* | *Listeria* |
| PSU-VKH-LP019 | MH341451 | *Siphoviridae* | unclassified | *Listeria* |
| PSU-VKH-LP040 | MH341452 | *Siphoviridae* | unclassified | *Listeria* |
| PSU-VKH-LP041 | MH341453 | *Myoviridae* | unclassified | *Listeria* |
| vB_LmoS_188 | NC_028871.1 | *Siphoviridae* | unclassified | *Listeria* |

**Table S2.** BLASTn search results for LGI-2 of *L. monocytogenes* ST14 persistent clade C8 strain HC87. Only results with 100% percentage nucleotide identity are shown.

| **Strain** | **Species** | **Query Coverage** | **Lenght** | **Accession** |
| --- | --- | --- | --- | --- |
| AUSMDU00000235 | *Listeria monocytogenes* | 100 % | 3005026 | CP045970.1 |
| J1-220 | *Listeria monocytogenes* | 100 % | 3032269 | CP006046.4 |
| 20 | *Listeria monocytogenes* | 100 % | 3000197 | CP030803.1 |
| FDAARGOS_611 | *Enterococcus faecalis* | 100 % | 2845446 | CP041012.1 |
| FDAARGOS_778 | *Listeria monocytogenes* | 100 % | 2950983 | CP040988.1 |
| FDAARGOS_607 | *Listeria monocytogenes* | 100 % | 2992056 | CP041014.1 |
| Li 2108 | *Listeria monocytogenes* | 100 % | 3012854 | CP039751.1 |
| 110 | *Enterococcus faecalis* | 100 % | 2865654 | CP039752.1 |
| M13455 | *Listeria monocytogenes* | 100 % | 2950983 | CP031476.1 |
| ATCC 13932 | *Listeria monocytogenes* | 100 % | 2950984 | CP025219.1 |
| PIR00546 | *Listeria monocytogenes* | 100 % | 3031937 | CP025220.1 |
| CFSAN054108 | *Listeria monocytogenes* | 100 % | 2994410 | CP028333.1 |
| VIMVR081 | *Listeria monocytogenes* | 100 % | 3054454 | CP018148.1 |

**Table S3.** Genes positively or negatively associated (Bonferroni-corrected *p*<.05) with persistent clades among the 233 *L. monocytogenes* Lineage II isolates of this study.

| **Element** | **Protein Accession** | | **Annotation or domain** | **Odds ratio** | **Bonferroni p** |
| --- | --- | --- | --- | --- | --- |
| type VII secretion system | | WP_026750033 | ESAT-6 secretion machinery protein EssC | 7.11 | 1.5E-04 |
| type VII secretion system | | WP_026750032 | DUF5082 | 7.11 | 1.5E-04 |
| type VII secretion system | | WP_003731422 | hypothetical protein | 6.95 | 1.9E-04 |
| hypervariable hotspot 4 | | WP_061668643 | hypothetical protein | inf | 2.1E-07 |
| hypervariable hotspot 8 | | WP_003733794 | hypothetical protein | 4.60 | 5.8E-03 |
| hypervariable hotspot 8 | | WP_026749868 | immunity protein (Imm-NTF2 -domain) | 4.60 | 5.8E-03 |
| hypervariable hotspot 8 | | WP_003721523 | DUF4274 | 4.49 | 1.1E-02 |
| IS3-like mobile element | | EAG2033583 | SAM-dependent methyltransferase, partial | 6.14 | 8.3E-05 |
| phage | | YP_009907771 | hypothetical protein | 8.33 | 6.7E-07 |
| phage | | YP_009044813 | hypothetical protein | 6.23 | 4.3E-05 |
|  | | WP_003722049 | 1,4-beta-N-acetylmuramoylhydrolase | 10.01 | 4.6E-04 |
|  | | WP_003722051 | ribonuclease BN | 7.21 | 4.3E-06 |
|  | | WP_009933017 | hypothetical protein | 7.73 | 2.1E-02 |
|  | | WP_026749741 | biofilm-associated protein BapL | 7.11 | 1.5E-04 |
|  | | WP_010989981 | hypothetical protein | 4.33 | 4.9E-02 |
| CRISPR-cas IIA | | WP_014601172 | CRISPR-associated endonuclease Cas9 | 0.00 | 3.2E-03 |
| CRISPR-cas IIA | | WP_014601170 | CRISPR-associated protein Csn2 | 0.00 | 6.7E-04 |
| CRISPR-cas IIA | | WP_003723648 | CRISPR-associated endoribonuclease Cas2 | 0.00 | 6.7E-04 |
| CRISPR-cas IIA | | WP_009925355 | CRISPR-associated protein Cas4 | 0.00 | 2.8E-14 |
| CRISPR-cas IIA | | WP_014601171 | CRISPR-associated endonuclease Cas1 | 0.00 | 6.7E-04 |
| type VII secretion system | | WP_05431447 | ESAT-6 secretion machinery protein | 0.09 | 4.9E-08 |
| type VII secretion system | | WP_009924118 | DUF4176 | 0.09 | 4.9E-08 |
| type VII secretion system | | WP_003724890 | hypothetical protein | 0.09 | 4.9E-08 |
| type VII secretion system | | WP_009924116 | hypothetical protein | 0.09 | 4.9E-08 |
| type VII secretion system | | WP_070275768 | hypothetical protein | 0.00 | 3.9E-08 |
| type VII secretion system | | WP_031674920 | toxin B | 0.00 | 3.9E-08 |
| hypervariable hotspot 8 | | WP_012951470 | hypothetical protein | 0.22 | 5.8E-03 |
| hypervariable hotspot 8 | | WP_009931454 | hypothetical protein | 0.21 | 5.0E-03 |
| hypervariable hotspot 8 | | WP_014931350 | DUF3130 | 0.17 | 1.6E-04 |
| hypervariable hotspot 8 | | ADB67935 | VOC family protein | 0.05 | 6.7E-10 |
| hypervariable hotspot 8 | | WP_052672896 | chromosome segregation ATPase | 0.00 | 2.8E-05 |
| hypervariable hotspot 8 | | WP_045131475 | hypothetical protein | 0.00 | 2.8E-05 |
| hypervariable hotspot 8 | | WP_031664941 | recombination and strand exchange inhibitor | 0.00 | 5.6E-06 |
| hypervariable hotspot 8 | | WP_045131476 | hypothetical protein | 0.00 | 2.8E-05 |
| hypervariable hotspot 8 | | WP_014600726 | AraC family transcriptional regulator | 0.00 | 4.0E-11 |
| LmoJ3 | | WP_061661643 | DNA cytosine methyltransferase | 0.00 | 2.8E-05 |
| LmoJ3 | | WP_069000933 | NgoFVII family restriction endonuclease | 0.00 | 2.8E-05 |
| LmoJ3 | | WP_069000932 | XRE-family transcriptional regulator | 0.00 | 2.8E-05 |
| phage | | YP_009907759 | hypothetical protein | 0.13 | 2.1E-02 |
| phage | | YP_009044855 | hypothetical protein | 0.07 | 2.7E-02 |
| phage | | WP_003725093 | hypothetical protein | 0.05 | 2.9E-02 |
| phage | | WP_003725092 | KTSC domain | 0.05 | 2.9E-02 |
| phage | | WP_009917698 | hypothetical protein | 0.04 | 2.1E-07 |
| phage | | NP_463476 | phage_Gp15 domain | 0.03 | 6.0E-03 |
| phage | | WP_014930203 | DUF1642 | 0.02 | 1.4E-03 |
| phage | | YP_001468391 | phage_GP20 | 0.02 | 3.0E-04 |
| phage | | NP_463533 | gp68 | 0.02 | 1.3E-05 |
| phage | | WP_020830757 | hypothetical protein | 0.00 | 6.7E-04 |
| phage | | NP_463467 | major capsid protein | 0.00 | 2.8E-05 |
| phage | | WP_003731437 | XRE family transcriptional regulator | 0.00 | 1.5E-02 |
| phage | | WP_014601102 | XRE family transcriptional regulator | 0.00 | 1.5E-02 |
| phage | | YP_001468393 | gp7 | 0.00 | 6.7E-04 |
| phage | | YP_009907764 | YozE_SAM_like -domain | 0.00 | 2.8E-05 |
| phage | | YP_009907767 | hypothetical protein | 0.00 | 1.5E-02 |
| phage | | NP_463462 | putative terminase small subunit | 0.00 | 2.8E-05 |
| phage | | YP_009907763 | XRE-family transcriptional regulator | 0.00 | 3.2E-03 |
| phage | | WP_097528476 | hypothetical protein | 0.00 | 3.2E-03 |
| phage | | YP_008126752 | Sipho_Gp157 | 0.00 | 3.2E-03 |
| phage | | YP_009907755 | GP47 protein | 0.00 | 1.5E-02 |
| phage | | WP_003733717 | helix-turn-helix domain-containing protein | 0.00 | 1.5E-02 |
|  | | WP_054314211 | cell wall anchor protein | 0.10 | 9.6E-07 |
